# KSHV Topologically Associating Domains in Latent and Reactivated Viral Chromatin

**DOI:** 10.1101/2022.04.05.487244

**Authors:** Mel Campbell, Chanikarn Chantarasrivong, Yuichi Yanagihashi, Tomoki Inagaki, Ryan R. Davis, Kazushi Nakano, Ashish Kumar, Clifford G. Tepper, Yoshihiro Izumiya

**Author notes:** Correspondence and request for materials should be addressed to Y.I. Address: UCDMC Research III Room 2200B, 4645 2^nd^ Avenue, Sacramento CA 95817, E. mail. Equal contribution (Mel Campbell, Chanikarn Chantarasrivong).

## Abstract

Eukaryotic genomes are structurally organized via the formation of multiple loops that create gene expression regulatory units called topologically associating domains (TADs). Here we revealed the KSHV TAD structure at 500 base pair resolution and constructed a 3D KSHV genomic structural model with 2kb binning. The latent KSHV genome formed very similar genomic architectures in three different naturally infected PEL cell lines and in an experimentally infected epithelial cell line. The majority of the TAD boundaries were occupied by CTCF and SMC1, and the KSHV transactivator was recruited to these sites during reactivation. Triggering KSHV gene expression decreased pre-wired genomic loops within the regulatory unit, while contacts extending outside of regulatory borders increased, leading to formation of a larger regulatory unit with a shift from repressive to active compartments (B to A). The 3D genomic structural model proposes that the immediate-early promoter region is localized on the periphery of the 3D viral genome during latency, while highly inducible non-coding RNA regions moved toward the inner space of the structure, resembling the configuration of a “bird cage” during reactivation. The compartment-like properties of viral episomal chromatin structure and its reorganization during the transition from latency may help coordinate viral gene transcription.

**Importance:** The 3D architecture of chromatin allows for efficient arrangement, expression, and replication of genetic material. The genomes of all organisms studied to date have been found to be organized through some form of tiered domain structures. However, the architectural framework of the genomes of large double-stranded DNA viruses such as the herpesvirus family has not been reported. Prior studies with Kaposi’s sarcoma-associated herpesvirus (KSHV) have indicated that the viral chromatin shares many biological properties exhibited by the host cell genome, essentially behaving as a mini human chromosome. Thus, we hypothesized that the KSHV genome may be organized in a similar manner. In this report, we describe the domain structure of the latent and lytic KSHV genome at 500 base pair resolution and present a 3D genomic structural model for KSHV under each condition. These results add new insights into the complex regulation of the viral lifecycle.

## Introduction

Kaposi’s sarcoma-associated herpesvirus (KSHV), also known as human herpesvirus-8 (HHV8), is a member of the gammaherpesvirus family of double-stranded DNA viruses. The virus is strongly associated with Kaposi’s sarcoma (KS), an endothelial cell-derived tumor, and two rare lymphoproliferative disorders, multicentric Castleman’s disease (MCD), and primary effusion lymphoma (PEL). KSHV exhibits a broad host range and can infect a variety of cell types *in vitro*, including B lymphocytic cells, renal-derived cells, and human gingival epithelial cells (1-3). KSHV DNA exists in the virion as a linear duplex of ∼140 kb of coding sequence which encodes approximately 80 genes or open-reading frames (ORFs) that are expressed in a highly-coordinated manner during latency or lytic replication (4, 5). The viral coding sequences are flanked on either side by tandem terminal repeats of highly GC-rich noncoding sequences giving rise to a genome of ∼160-170 kb (6, 7). Following infection, the chromatin-free viral genome circularizes, is rapidly chromatinized (8) and maintained and replicated in the nucleus of the host cell as monomeric episomes. A crucial element of the KSHV lifecycle is the reactivation of the virus from a latent (dormant) state into the lytic replicative cycle in which viral genomic DNA is replicated, viral particles are produced, and then virions are released along with host cell lysis.

A great deal of effort has focused on defining the epigenetic mechanisms that regulate latency, reactivation, and commandeering of the host cell by KSHV (for reviews, see (9-11). Although a single viral protein, K-Rta (ORF50), triggers the onset of lytic replication by the transcriptional activation of lytic genes, the exact details of how K-Rta functions as this master regulator have been enigmatic. We have previously shown that interplay between KSHV-encoded transcription factors and host cell-encoded epigenetic regulators (JMJD2A demethylase, EZH2 methyltransferase) are key mechanisms in controlling KSHV reactivation (8, 12). In addition, using Capture Hi-C (chromosome conformation capture analyses with deep sequencing) analyses of the KSHV genome, we demonstrated that K-Rta not only activates individual lytic genes by binding to specific regulatory elements along the KSHV genome, but that it also performs a higher-order coordination of the process by mediating three-dimensional (3D) conformational changes in the architecture of the KSHV chromosome (13). In essence, “chromatin loops” are formed via K-Rta promoting contact between distant genomic regions containing key regulatory sequences.

Advances in chromosome conformation capture (3C)-based studies have uncovered the existence of spatially insulated genomic regions that are now considered the invariant building blocks of chromosomes (14-17). When initially reported in 2012, these ∼100 kb-1 Mb regions were referred to as topological domains (14) or topologically associating domains (TADs) (15). These domains are defined by the preferential interaction of loci located within a given TAD and a relative (∼2-fold) depletion of interactions between loci located in different TADs (16). In the era of 3C-based methods, TADs were originally described in mammalian cells (14, 15) using high-throughput chromosome conformation capture (Hi-C) (18) or chromosome conformation capture carbon-copy (5C) (19), and in *Drosophila* (17) chromosomes using Hi-C. However, TADs have also been found in Zebrafish (20), *Caenorhabditis elegans* (21), *Saccharomyces cerevisiae* (22) and *Schizosaccharomyces pombe* (23). In addition, analysis of nucleoids of *Caulobacter crescentus* (24) and *Bacillus subtilis* (25, 26) have described genomic spatial domains termed Chromosomal Interaction Domains (CIDs) which mimic TADs in terms of preferential interaction properties. Together, these results suggest that TAD-like domains, although differing in size among the genomes of the various species analyzed (27), may be a universal feature of both prokaryotic and eukaryotic genomes.

The existence of TAD-like structures within the genomes of large double-stranded DNA viruses such as herpesvirus family has not been reported. For KSHV there are numerous reports concerning the role of the structural maintenance of chromosomes (SMC) cohesin complex and CCCTC-binding factor (CTCF) in control of viral latency and reactivation (28-36). As both cohesin and CTCF have been linked to chromatin looping formation in mammalian cells (14, 37-39), this suggests the plausible existence of TAD-like structures within KSHV genomes. Moreover, previous 3C (33) and capture Hi-C (13) analyses have documented the presence of chromatin looping within the latent and lytic KSHV genomic regions. Together with KSHV epigenomic mapping data (8, 12, 40-43), this suggests that the chromatinized KSHV genome shares many biological properties exhibited by the host genome, behaving as a mini human chromosome and, as such, may be organized in a TAD-like manner. In this report, we describe the TAD-like structures of the latent and lytic KSHV genome at 500 base pair (bp) resolution and present a 3D genomic structure model for the KSHV under each condition.

## Materials and Methods

### Capture Hi-C

KSHV Capture Hi-C (CHi-C) analysis was performed using a robust *in situ* CHi-C protocol with kitted reagents from Arima Genomics (San Diego, CA) based on well-accepted methods described for *in situ* Hi-C (38, 44), CHI-C (45-47), and as described previously (13). Briefly, cells were crosslinked with 2% formaldehyde, lysed, and genomic DNA digested with a cocktail of 4-cutter restriction endonucleases by incubation for 30 minutes at 37°C. The 5’-overhangs were then filled in and “marked” with biotin by incorporating biotinylated dATP (biotin-14-dATP) with Klenow fragment of DNA polymerase I (incubation for 45 minutes at 25°C). Proximity intramolecular ligation of the blunt-ended fragments was then performed with T4 DNA ligase (incubation for 15 minutes at 25°C). The formaldehyde crosslinks were reversed, and the ligated, chimeric DNA products were purified with Agencourt AMPure XP paramagnetic beads (Beckman Coulter). The DNA was then sheared to an average length of 400 bp using a Covaris E220 Focused-ultrasonicator (Covaris, Inc.) and fragments size-selected with AMPure XP beads to achieve a size distribution of 200-600 bp. The biotin-marked ligation products were then enriched by affinity capture with streptavidin magnetic beads (DynaBeads MyOne Streptavidin C1; Invitrogen, Thermo Fisher Scientific). Subsequently, libraries were prepared from the bound ligation products with the Kapa HyperPrep Kit with Library Amplification Module (Roche) using an on-bead modification to the standard protocol for end repair, dA-tailing, and ligation of Illumina TruSeq sequencing adaptors.

The KSHV CHi-C library was then prepared by target enrichment of the libraries for KSHV genomic content by solution hybridization with a custom-designed KSHV genomic capture probe library (xGen Lockdown Probes; Integrated DNA Technologies, Inc., Coralville, IA) as previously described (13). Briefly, libraries (500 ng) were hybridized with the KSHV genomic capture probe pool (3 pmol) in a mixture containing xGen 1X Hybridization Buffer, Cot-1 (5 μg), and xGen Universal Blocking Oligos for 4 hours at 65°C. The hybridized targets were then captured with streptavidin beads (DynaBeads MyOne Streptavidin C1; Thermo Fisher Scientific) by incubation for 45 minutes at 65°C. Unbound DNA was removed by a series of high-stringency (65°C) and low-stringency (room temperature) washes. The KSHV genome-enriched CHi-C library DNA was eluted and PCR enrichment (12 cycles) performed with high-fidelity KAPA HiFi HotStart DNA Polymerase (Kapa Biosystems, Inc., Wilmington, MA). Libraries were multiplex sequencing (2 × 150 bp, paired-end, ∼30 million mapped reads/mate pairs per sample) on an Illumina Hiseq 4000 sequencing system (Supplement Figure S1).

### Hi-C data pre-processing

Sequence alignment and quality check of reads were performed using HiC-Pro 2.11.1 pipeline (48); each read-end was aligned to the human GRCh37 (hg19) and KSHV (NC_009333.1) reference genomes. Quality reports showed percentage duplicates at less than 15% while >95% di-tags were valid and aligned to restriction fragments (S-Figure 1). The duplicates were removed and reads mapping only to the KSHV genome were extracted by using Pysam 0.14.1.

### Genomic domain analysis

The KSHV genome was analyzed and visualized for frequencies of chimeric genomic reads, and finally modeled for 3D Structure using TADbit (49). The KSHV mapped reads were filtered for only uniquely mapped reads pairs based on the intersection of each read-end. The reads that were self-ligations, dangling-end, error, extra dangling-end, too short, too large, duplicated, and random breaks were filtered out to provide valid reads. The valid reads were stored as matrices and binned with resolution of 500 bp (2 kb for 3D modelling); the bins with more than 1000 counts and at least 75% of cells with no-zero counts were used in the next steps. Iterative Correction and Eigenvector decomposition (ICE) normalization was used to treat the data with iteration=100. Bins were identified as in compartment A or B by calculating eigenvectors. TADs/TAD borders were called, and insulation score/border strength were calculated using the TADbit computational framework (49), and visualized as contact maps and aligning plots.

### KSHV 3D chromosome structure modelling

Matrix Modeling Potential (MMP) score (50), ranging from 0 to 1, was used to identify if the interaction matrices from the previous step (i.e., iterative modeling with TADbit), have potential for modeling. The higher the MMP scores are, the more likely the 3D model will be accurate. In this study, MMP scores of latency and reactivation models were 0.9541 and 0.9023, respectively. Monte Carlo optimization method was used to optimized parameters from both models. The parameters included maximal distance associated between two interacting particles, particles attraction, particles repulsion, and contact distance of particles. Sets of models were produced from the possible combinations of those parameters. Contact maps for each set of models were built, then compared with the Hi-C interaction by averages of Spearman correlation coefficients. The models with higher correlation coefficients represented the original data and were chosen to be visualized and plotted by DNA density. The correlations of the latency model and reactivation models were 0.8706 and 0.8782, respectively. Finally, three-dimensional visualization of KSHV molecular modeling were performed with the UCSF Chimera package (51).

### Cleavage Under Targets and Release Using Nuclease (CUT&RUN)

CUT&RUN (52) was performed essentially by following the online protocol established by Dr. Henikoff’s lab with several modifications. Cells were washed with PBS and wash buffer [(20 mM HEPES-KOH pH 7.5, 150 mM NaCl, 0.5 mM Spermidine (Sigma, S2626) and proteinase inhibitor (Roche)]. After removing the wash buffer, cells were captured on magnetic ConA beads (Polysciences, PA, USA) in the presence of CaCl_2_. Beads/cells complexes were washed 3 times with digitonin wash buffer (0.02% digitonin, 20 mM HEPES-KOH pH 7.5, 150 mM NaCl, 0.5 mM Spermidine and 1x proteinase inhibitor), aliquoted, and incubated with specific antibodies in 250 μL volume. The antibodies used in this study were: rabbit polyclonal anti-SMC1 (Cell Signaling, #4802S; 1:100), mouse anti-RNA polymerase II (Millipore, clone CTD4H8; 1:100), and rabbit monoclonal anti-CTCF (Cell Signaling, clone D31H2, #3418S; 1:100). After incubation, unbound antibody was removed by washing with digitonin wash buffer 3 times. Beads were then incubated with recombinant pAG-MNase, which was purified from *E*.*coli* (S-Figure 1), in 250 μl digitonin wash buffer at 1.0 μg/mL final concentration for one hour at 4°C with rotation. Unbound pAG-MNase was removed by washing with digitonin wash buffer 3 times. Pre-chilled 2 mM CaCl_2_ containing digitonin wash buffer (200 μL) was added to beads and incubated on ice for 30 min. The pAG-MNase digestion was halted by the addition of 200 μl 2× STOP solution (340 mM NaCl, 20 mM EDTA, 4 mM EGTA, 50 μg/ml RNase A, 50 μg/ml glycogen). The beads were incubated with shaking at 37°C for 10 min in a tube shaker at 500 rpm to release digested DNA fragments from the insoluble nuclear chromatin. The supernatant was collected after centrifugation (16,000 x g for 5 min at 4°C) and placed on a magnetic stand. DNA was extracted using the NucleoSpin kit (Takara Bio, Kusatsu, Shiga, Japan). Sequencing libraries were then prepared from 3 ng of CUT&RUN DNA with the Kapa HyperPrep Kit (Roche) according to the manufacturer’s standard protocol. Libraries were multiplex sequenced (2 × 150bp, paired-end) on an Illumina HiSeq 4000 sequencing system to yield ∼15 million mapped reads per sample. When necessary, *E. coli* genomic DNAs carried over in purified pAG-MNase was used to normalize sequence reads as described previously (52).

### Genomic data analysis

FASTQ files for the capture Hi-C experiments were processed through the HiCUP (v0.7.4) pipeline (53) using the KSHV (NC_009333.1) genome. Valid interaction products called by HiCUP were converted into Juicebox (54) input format (.hic file), which stores the normalized and un-normalized contact matrices as a highly compressed binary file, by using a series a scripts provided by HiCUP (hicup2homer) and HOMER (makeTagDirectory and tagDir2hicFile) (55). Juicebox was utilized to facilitate adjustments of resolution and normalization, intensity scaling, zooming, and addition of annotation tracks.

CUT&RUN sequence reads were aligned to the human hg38 reference genome assembly and reference KSHV genome sequence with Bowtie2 v2.3.5.1 (56) and/or HISAT2 v2.1.0 (57). MACS2 (Model-based Analysis of ChIP-Seq) v2.1.1.1.20160309 was used for detecting peaks (58) following the guidelines in the developer’s manual. The following publicly available data sets from previous studies (59, 60) were also used to overlay binding sites or histone modification sites; KSHV K-Rta ChIP-seq (GSE123897), H3K27Ac, H3K4me3, and H3K27me3 (GSE163695). Peaks and read alignments were visualized using the Integrated Genome Browser (IGB) (61). Heatmaps and average profile plots were drawn from bed files created by MACS and bam files using the R package, ngsplot v2.63 (62).

### Cell Culture, bacmid transfection, and selection

TREx-F3-H3-K-Rta BCBL-1 cells (TREx-BCBL-1) cells were maintained and reactivated as previously described (13). BC-1 and BC-3 cells were maintained in RPMI 1640 supplemented with 15% FBS, 2mM glutamine, and 1% penicillin-streptomycin. BC-1 and BC-3 cell lines were obtained from the ATCC (Manassas, Va). *i*SLK cells that were infected with BAC16 HA-ORF57 Wt virus, were maintained in DMEM supplemented with 10% FBS, 1% penicillin-streptomycin-L-glutamine solution, 1 mg/ml hygromycin B, and 250 μg/ml G418. *i*SLK cells were obtained from Dr. Don Ganem (Novartis Institute for Biomedical Research) and were maintained in DMEM supplemented with 10% FBS, 1% penicillin-streptomycin-L-glutamine solution and 400 μg/ml hygromycin and 250 μg/ml G418 and 10μg/ml puromycin.

## Results

### KSHV topologically associating domains

In our previous study, we demonstrated that the KSHV genome forms organized DNA loops in infected cell nuclei and the frequency of these loops increased during reactivation, especially near K-Rta binding sites (13, 63). In order to understand the molecular details of loop formation, we have performed additional Capture Hi-C experiments to augment the results from our earlier publication. Previously, our experimental workflow utilized an in-house 3C procedure based on Gavrilov et al. (64) with a single restriction enzyme digestion prior to library capture with KSHV lockdown probes (13). In the current study, a commercially available Hi-C kit (Arima Genomics) was used in conjunction with our probe library. In this procedure, the chromatin digestion step is performed *in situ* and employs a restriction enzyme cocktail coupled with fill-in reactions to incorporate a biotinylated deoxynucleotide to mark the digested ends which facilitates the enrichment of the DNA ligation products (i.e., consisting of the spatially proximal digested ends) prior to the KSHV lockdown probe capture step. The restriction enzyme digestion generated fragment sizes with an average length of 190 bp +/- 210 bp. The largest fragment is 2,225 bp, which is located near the LANA coding region, while the shortest fragment is 4 bp. The distribution of fragment sizes is shown in S-Figure 2B. A total of 3,565,321 valid read pairs were uniquely mapped on the KSHV genome in non-reactivated BCBL-1 cells, while 5,967,349 valid read pairs were mapped in the reactivated sample. Using two capture/enrichment steps, we further increased the number of sequence reads corresponding to valid Hi-C di-tags by approximately 1,000-fold over our previous studies and prepared KSHV genomic interaction maps at 500 bp resolution (Figure 1, S-Figure 2). Because topologically associating domains often refer to sizes of 100 kb to 1 Mb genomic domains (14), we decided to use the term of transcription regulatory domain (TRD).

**Figure 1.**
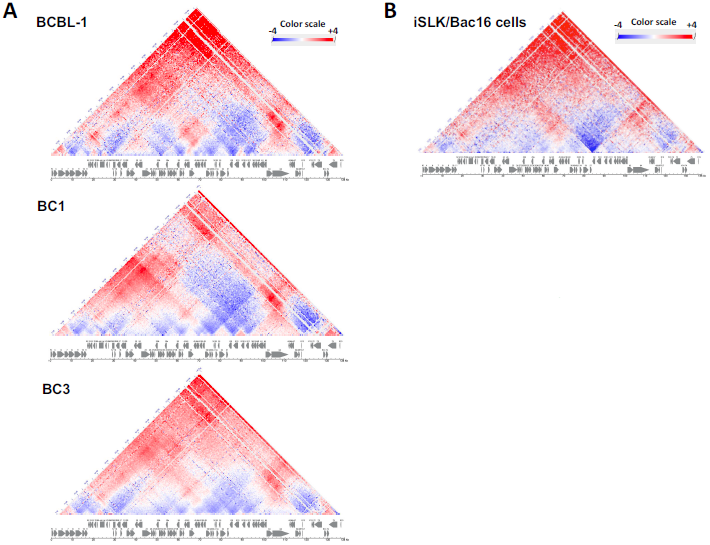
Normalized frequencies of genomic contact map during latency at 500 bp resolution. **(A) KSHV genome contact maps in latent PEL cells.** Genomic contacts detected at more than expected frequencies are depicted in red, while underrepresented contacts between loci are presented in blue. Contact maps are shown for latent BCBL-1, BC1 and BC-3 cells. **(B)** Genomic contacts detected in latent *i*SLK/BAC16 cells are depicted as described in (A). A color range scale is listed and KSHV ORF maps are shown beneath each contact map.

The KSHV episomes present in three naturally infected PEL cell lines (BCBL-1, BC1 and BC3) all exhibited very similar TRD structures (Figure 1). In addition, we also determined KSHV TRD structure in experimentally-infected *i*SLK/Bac16 cells. The results demonstrated very similar TRD structures to those of the naturally-infected cells. Our TRD map also agreed with previous 3C studies that positioned the LANA promoter region forming genomic loops with K12 loci at higher frequencies (31). By overlaying previously determined histone modification and transcription factors binding sites, we next examined the association with TRD formation (Figure 2). Although the genomic domains are not defined by chromatin state, sites of histone modification aligned very well with TRD formation and were organized with genomic domains with active histone marks (H3K4me3, H3K27Ac). In fact, the genomic regions with active histone marks seem to be physically neighboring each other in the 3D structure, which is seen by higher frequencies of genomic ligations than the theoretically-expected ligated sequence reads (shown in red color, where two H3K4me3 marked genomic regions were ligated more frequently than theoretically-expected frequency), and these features were common in all cell lines tested. The results also demonstrated that TRDs could also be distinctly separated based on the reactivation kinetics class of KSHV gene expression. For instance, late gene clusters were localized in distinct TRDs (within separated triangles). Consistent with previous reports, CTCF localizes to the junctions of the majority of TRDs, and CTCF indeed colocalized with SMC1 on the KSHV genomes (Figure 2B, C). In addition, superimposing K-Rta recruitment sites that were reported previously (59) further showed that K-Rta recruitment sites (Figure 2C black line) localized adjacent to CTCF/SMC1 binding sites (Figure 2C, red and blue lines), and the genomic regions harbored poised/stalled RNAPII (Figure 2C shadow). The results indicate that the KSHV episome is structured to allow the K-Rta complex to access to other genomic regions through genomic looping, which may serve to maximize K-Rta transactivation function. The NGS plot further confirmed that K-Rta binding sites are closely localized to CTCF and RNAPII binding sites (Figure 2D).

**Figure 2.**
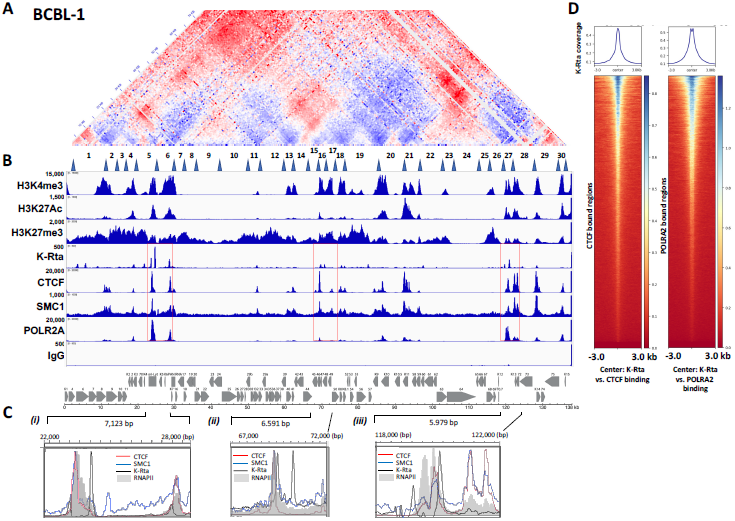
Superimposition of select histone modifications and protein factor binding sites on genomic contacts of the latent KSHV genome. (A) The BCBL-1 contact map from Figure 1A is shown. Latent TRDs numbered 1-30 with their borders (blue triangles) (from Fig. 3A/B) are listed below the contact map. (B). Alignment of histone modifications H3K4Me3, H3K27Ac, H3K27me3, and binding sites of K-Rta, CTCF, SMC1, POLR2A are shown. Control IgG for background control is also shown. The sequence reads were mapped to the KSHV genome. IGV snapshots and a KSHV genome map are shown. Numbers on the left-hand side of each track denotes the height of the peak (e.g., read depth). **(C). Zoomed view of CTCF, SMC1, K-Rta, and RNAPII (POLR2A) enrichment at select regions of the KSHV genome** (i) K4-PAN RNA, (ii) ORFs 45-50 and (iii) repeat region. **(D). Positional correlations among K-Rta, CTCF, and RNAPII recruitment sites**. Density plot showing average K-Rta ChIP-seq signals within ±3 kb regions around the center of CTCF or PORLA2 peaks. Heatmap shows K-Rta signals on CTCF or POLRA2 CUT&RUN peaks. *y*-axis is ranked according to CTCF or PORLA2 enrichment in descending order.

### Regulation of TAD structure during reactivation

Having defined the KSHV genomic TRD structure in the condition that the majority of episomes were in during the latent state, we next examined changes in TRDs induced during the early stages of reactivation. To avoid complications with ongoing DNA replication, we examined a time point (i.e., 24 hrs post-reactivation) at which there was an only minimal increase in viral DNA copies, as measured by qPCR (60). Capture Hi-C analyses were performed with TREx-BCBL-1 cells at 0 and 24 hours post-reactivation using an induction protocol consisting of a combination of TPA and doxycycline. The experiments were performed in triplicate. Using the TAD caller program TADbit for visualization of hierarchical genomic domains, the formation of larger TRDs during reactivation was observed when compared with TRD structures observed during latency, and presumably proceeded through the fusion of neighboring TRDs via assembly of active transcription sites, [Figure 3A (a, b), (63)]. The results also showed that pre-existing genomic loops in latent cells (Figure 1 in red) were largely disrupted during reactivation, as indicated by a marked decrease in the frequency of their interactions within individual TRDs (Figure 3B, visualized as blue color). While pre-existing TRDs were disrupted, genomic contacts that spanned outside of the previously existing TRD borders were increased (Figure 3B, visualized as red color). Reactivation was also characterized by new interactions of the K-Rta promoter region formed with the late gene clusters (ORF63-68), while pre-formed, latency-associated contacts with active compartments, such as E gene clusters (ORF6-ORF11) were decreased (Figure 3B).

**Figure 3.**
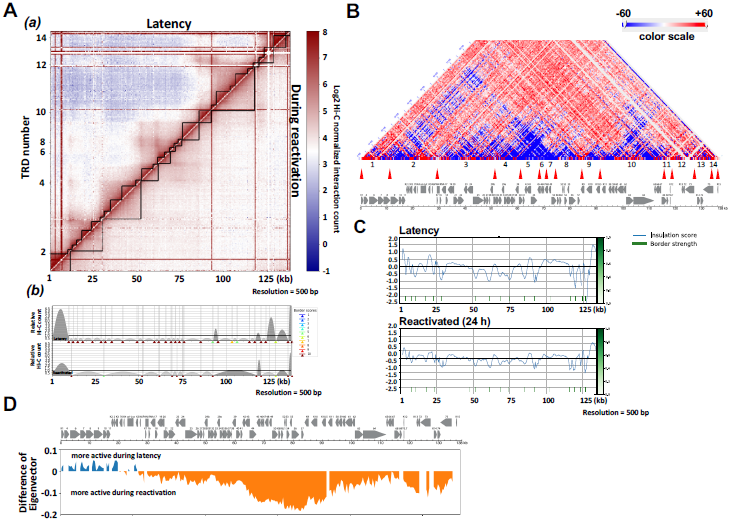
Dynamics of genomic contacts by reactivation in TREx-BCBL-1 cells. **(A) Interaction Frequency Matrix**. (a) TRDs were calculated and visualized with TADbit, and genomic domains defined by frequencies of DNA contacts are marked with black solid lines. (b) TAD border alignment during latency (upper) and during reactivation (lower) are shown. Dark and light grey arches indicate TRDs with higher and lower than expected intra-TRD interactions, respectively. TADbit border robustness (from 1 to 10) is identified by a color gradient from blue to red. **(B) Contact Heatmap: BCBL-1 reactivation vs. latency**. Contacts are color coded as described in Figure 1 (red, increased contacts; blue decreased contacts). TRDs called during reactivation (Fig. 3A/B) numbered 1-14 with their borders (red triangles) are listed below the contact map. KSHV ORF map is included in the bottom panel. **(C) TAD border analyses**. (a)Insulation score (border strength) is plotted. Reactivation reduced insulation and border strength to create larger transcription units. **(D)** Compartment analysis. 500 bp bins were identified as in compartment A or B by calculating eigenvectors.

Next, insulation scores were calculated to reveal changes in TRDs before and after reactivation. A lower score suggests a higher insulation effect, which is indicative of the position of the boundary. The border strength at the majority of genomic regions was decreased in reactivating samples (Figure 3C), suggesting more freely interaction with other genomic domains when viral lytic gene expression was triggered. As expected, compartment analyses showed shifts from repressive to active compartments (B to A) during reactivation [Figure 3D]. Interestingly, the analyses also showed that E gene clusters encompassing sequences from K1 to PAN RNA coding regions (genomic locations bin 1-60 [1-30 kb region]) showed an active compartment structure in latent chromatin. This result is somewhat unexpected since these genomic loci largely encode lytic genes. While LANA, two Ori-Lyts regions, K10.5/11, and the PAN RNA coding region did not change a specific compartment structure between that observed during latency and after triggering reactivation, these genomic regions are narrowly insulated with higher CTCF/SMC1 peaks (Figure 2B).

### KSHV 3D genomic structure, a bird cage model

With 3,565,321 (latent) and 5,967,349 (lytic) valid Hi-C di-tags (i.e., genomic contacts) covering the ∼140 kb KSHV genome, we next constructed a theoretical 3D KSHV genomic structural model utilizing TADbit, which was then visualized with the UCSF Chimera program (51). The model proposes that the immediate-early promoter region (e.g., K-Rta) is localized on the periphery of the 3D viral genome during latency, which is likely to be more accessible to the nuclear environment (Figure 4, Supplemental material movie); while the highly inducible long non-coding RNA region (i.e., PAN RNA) moved toward the inner space of the structure in reactivating episomes. During the transition from latency to the lytic cycle, the overall 3D genomic structures were squeezed into a doughnut-like configuration from a spherical shape. This arrangement positions the transcriptionally active regions to be more closely neighboring to other genomic loci in three-dimensional space, and their conformation change approximates a “bird cage” (Figure 4). These changes were also consistent with disruption of pre-existing TADs and the increase in genomic loops extending outside of TAD domains as described in Figure 3A. Taken together, we propose that the KSHV 3D genomic structure is designed to maximize the effects of K-Rta complex recruitment and may also facilitate re-utilization of active RNAPII complexes.

**Figure 4.**
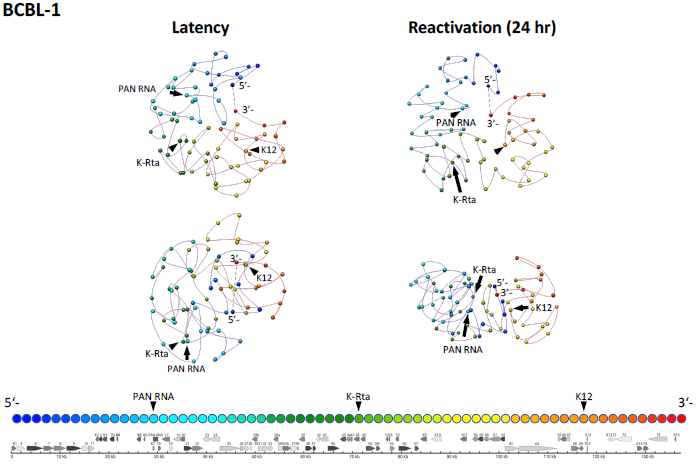
KSHV 3D genomic structure modeling. Frequencies of ligated fragments were used to calculate distances between fragments at 2 kb resolution. The average of 8 different models were used to draw the figures. KSHV 3D genomic structure before (left) and after (right) triggering reactivation are shown. Color gradation associated with KSHV genomic position are depicted at the bottom of figures. The movies of the 3D figures including individual 8 models in shadow are presented in Supplemental Figure S2. The dotted line represents the terminal repeat (TR) region. Each sphere marks 2 kb of linear genomic sequence.

## Discussion

In this report, we have empirically mapped the genomic contacts within the KSHV genome during latency and at an early stage of reactivation. This mapping, combined with bioinformatic tools, has allowed us to uncover TAD-like structures of the KSHV genome during each condition. Using a workflow consisting of HiC-Pro and TADbit, we have constructed a structural model which allows visualization of the overall conformational changes that occur in the viral genome during this time period. With the analyses, similar contact maps were obtained among three PEL cell lines and one laboratory infected cell line (*i*SLK BAC16-Wt). This overall similarity suggests that the prominent viral genome contacts established during latency appear to be conserved among different KSHV infected cell lines and are stable properties within the infected cell population.

Previous studies have implicated CTCF sites as being important for several KSHV genomic loops that involve the latency control region and either the K12 or ORF50 locus (31). Inspection of the contact heat maps aligned well with the CTCF/SMC1 recruitment sites at the corners of TRDs, supporting the notion that CTCF/SMC1 plays a major role in defining TRD domains. However, the smaller TRDs also show a lack of CTCF/SMC1 (Figure 2), possibly reflecting that enriched contacts are not readily visible due to the transient nature of extrusion (65) or that other architectural protein factors could also play a role in structuring latent KSHV genomes. In this respect, KSHV latent genome structures resemble “ordinary domains” as previously defined by Rao et al. (38) to classify contact domains not flanked by CTCF binding sites and formed without involvement of CTCF/cohesin-mediated loop extrusion. As those smaller TADs are separated by transcription units (e.g., gene clusters with genes transcribed in the same direction and having similar kinetics), we speculate that RNA-binding proteins may also play a role in viral genomic structure (66). Although we have highlighted contacts between the ORF50 and K12 loci [(13), Figure 4] and a potential lncRNA from this region, it should be noted that it is uncertain which gene products originating from promoters in this region that might be influenced by genomic looping. This region harbors an origin of lytic replication and the lytic ALT lncRNA running antisense (67). Additional possible candidates are mRNAs encoding the Kaposin locus ORFs as well as a lytic miR-K12-10/12 pri-miRNA (4, 68).

During the shift from latency to lytic replication viral genomes exhibited a loss of local proximal intra-TRD contacts with the widespread onset of inter-TRD contacts (Fig. 3A(a)). Evidence exists for both dynamic (69-71) and stable (71-74) enhancer-promoter interactions during cellular responses to various stimuli or conditions. In different cell lines or across species TADs are generally conserved at the megabase scale (14, 39). However, when probed at higher levels of resolution, TADs may merge or be disrupted during changes in gene expression associated with a variety of cellular events such as development (75), lineage commitment (37), reprogramming (76), differentiation (77, 78), senescence (79) and possibly heat shock (80, 81). Together, the current study and our previous report (13), as well as findings from the Lieberman group (31) confirm the existence of both pre-wired and induced genomic loops and the dynamic nature of KSHV genomic contacts during the viral life cycle.

Insulation scores revealed the strength of borders on the KSHV genome. CTCF, SMC1, and K-Rta frequently bound at the borders, and these sites were decorated by H3K4me3 or H3K27Ac modifications. This observation suggested that K-Rta targets the boundaries established by CTCF and SMC1 with open chromatin structures. By targeting the boundaries, it is likely that K-Rta can recruit K-Rta-bound chromatin remodeling factors (82) to the multiple pre-assembled genomic regions for synchronous activation. In turn, Mediator complex, which is also a part of the K-Rta activation complex (82, 83), may help to establish active genomic hubs for future reactivation.

The border strength of most viral genomic regions decreased during lytic reactivation. This decrease suggests for greater accessibility of a given TRD to interact with other genomic elements, which is accompanied by induction of viral lytic gene transcription. However, the latency-associated loci remain relatively unchanged (Figure 3C). The latency genomic region is, in fact, localized in a small valley, which is well-protected by two strong peaks of CTCF and SMC1 in the KSHV genome (Figure 1B; genome coordinates 124-130 kbp). The result indicated that the majority of episomes in an infected cell within the given cell populations have similarly maintained CTCF and SMC1 sites. The result also suggests that these highly-utilized CTCF/SMC1 binding sites are likely to be essential for maintaining KSHV episomes in infected cells.

The dynamic nature of the bird cage model is consistent with results obtained using super-resolution imaging of mammalian cell chromatin structures where active histone marks coincided with less compact chromatin and exhibited a higher degree of co-localization with other active marks and RNAP II, while repressive marks coincide with densely packed chromatin and are spatially distant from active RNAP II (84). Wang et al. (85) also noted a decrease in chromatin fiber width when imaging was conducted using nuclei prepared from a quiescent state compared with nuclei undergoing active transcription. These data imply that in the quiescent state, chromatin fibers were more tightly packed together. This was contrasted by chromatin fibers imaged in the actively transcribing state, which appeared to be loosely entangled with each other and form a relatively thick structure.

We do recognize several caveats in our analysis, which is based on calculations assuming linear chromosomes while the KSHV genome is circular, a topology which has implications for interpretation of contact frequencies. Terminal repeats (TR) that occupy approximately one sixth of KSHV genome (e.g. 28 kb [0.8 kb x 35 copies]) were also not considered in our analyses due to difficulties in mapping sequence reads from this region. Further studies are therefore needed to confirm where and how a TR interacts with a unique region, which would have significant impact in KSHV reactivation through recruitment of the LANA protein complex (60). In addition, we employed a single TAD prediction tool for our analysis; one that allows for 3D modeling. However, it has been documented that TAD prediction algorithms may be very sensitive to data resolution and normalization resulting in calls that vary greatly between tools in terms of TAD number, size, and certain biological properties such as CTCF enrichment around boundaries (86, 87). At the same time, we do have confidence in the 3D structural model of the KSHV genome generated herein and the adequacy of the TADbit approach, based on its assessment conducted by simulations on artificially generated genomes, including a 1-Mb circular chromosome, and subsequent partial verification by fluorescence *in situ* hybridization imaging (50).

In summary, this study established 3D KSHV genomic structure models of episomes during latency and early lytic phase of reactivation and revealed the dynamics of TAD structural changes by transcription activation. The 3D genomic structures bestow another element to the effectiveness of herpesvirus gene regulation, a system which may provide an opportunity to further study TAD regulation through the intersection of viral and host proteins.

## Data Availability

The data sets supporting the conclusions of this study are available in the National Center for Biotechnology Information Gene Expression Omnibus (NCBI GEO) repository with the accession code GSE163695

## Acknowledgements

We would like to thank Drs. Kenichi Nakajima and Chie Izumiya for technical assistance; other members of the Izumiya lab for discussion and critical reading of the manuscript.

---

**Supplemental Figure S1**.

Preparation of pAG-MNase for CUT&RUN. Protein A or A&G domain fused MNase (pAG-MNAse) was expressed in *E*.*coli*, and purified with Ni-NTA beads (Invitrogen). Purified proteins were subjected to SDS-PAGE and stained with Coomassie blue. Recombinant protein sequence and purified proteins are shown. The pAG-MNase was used for CUT&RUN studies described in this manuscript.

**Supplemental Figure S2**.

**(A)** Representative valid sequence pairs in CHi-C. Overall sequence reads with invalid sequences were summarized in bar charts. One of three replicates of BCBL-1 samples for both before (top) and after triggering reactivation (bottom) are shown. **(B)** Predicted size distribution of KSHV genomic restriction fragments generated during CHi-C library preparation.

**Supplemental Figure S3 (Movies)**.

KSHV 3D genomic structure models for both latency and reactivated samples are shown. Gray shadow indicates each model and color string 5’ (blue) to 3’ (red) are generated based on average of 10 different models. Blue ball (PAN RNA region), Red ball (K-Rta promoter region), and Yellow ball (K12 genomic region) were marked for clarity.

